# A brief real-time fNIRS-informed neurofeedback training of the prefrontal cortex changes brain activity and connectivity during subsequent working memory challenge

**DOI:** 10.1101/2023.03.14.532684

**Authors:** Xi Yang, Yixu Zeng, Guojuan Jiao, Xianyang Gan, David Linden, Dennis Hernaus, Chaozhe Zhu, Keshuang Li, Dezhong Yao, Shuxia Yao, Yihan Jiang, Benjamin Becker

## Abstract

Working memory (WM) represents a building-block of higher cognitive functions and a wide range of mental disorders are associated with WM impairments. Initial studies have shown that several sessions of functional nearinfrared spectroscopy (fNIRS) informed real-time neurofeedback (NF) allow healthy individuals to volitionally increase activity in the dorsolateral prefrontal cortex (DLPFC), a region critically involved in WM. For the translation to therapeutic or neuroenhancement applications, however, it is critical to assess whether fNIRS-NF success transfers into neural and behavioral WM enhancement in the absence of feedback. We therefore combined single-session fNIRS-NF of the left DLPFC with a randomized sham-controlled design (N = 62 participants) and a subsequent WM challenge with concomitant functional MRI. Over four runs of fNIRS-NF, the left DLPFC NF training group demonstrated enhanced neural activity in this region, reflecting successful acquisition of neural selfregulation. During the subsequent WM challenge, we observed no evidence for performance differences between the training and the sham group. Importantly, however, examination of the fMRI data revealed that - compared to the sham group - the training group exhibited significantly increased regional activity in the bilateral DLPFC and decreased left DLPFC - left anterior insula functional connectivity during the WM challenge. Exploratory analyses revealed a negative association between DLPFC activity and WM reaction times in the NF group. Together, these findings indicate that healthy individuals can learn to volitionally increase left DLPFC activity in a single training session and that the training success translates into WM-related neural activation and connectivity changes in the absence of feedback. This renders fNIRS-NF as a promising and scalable WM intervention approach that could be applied to various mental disorders.

## 1. Introduction

Working memory (WM) conceptualizes a cognitive system that has been referred to as the “sketchpad of conscious thought”, given its essential role in all complex cognitive processes, ranging from problem-solving and decisionmaking to social cognition and language processing. Impairments in WM have been reported in a range of neurological and mental disorders, such as Alzheimer’s and Parkinson’s disease, schizophrenia, major depression, and substance use disorders (Lee and Park 2005; Forbes et al. 2009; Becker et al. 2010; Jahn 2013; Wang et al. 2015; Nikolin et al. 2021; Ramos and Machado 2021), and have been associated with impaired social and daily life functioning (Huang et al. 2014; Xu et al. 2022; Sun et al. 2023). WM thus represents a promising treatment target and initial studies have developed and evaluated behavioral and pharmacological interventions to enhance WM (Hernaus et al. 2017; Kaser et al. 2017; Teixeira-Santos et al. 2019; Zhao, Becker, et al. 2019; Afsaneh and Zarei Ghobadi 2022). During the last years brain stimulation techniques, such as transcranial direct current stimulation (tDCS) and transcranial magnetic stimulation (TMS) have been increasingly explored as potential strategies to improve WM capacity by directly targeting the brain systems that underlie WM. However, the efficacy of these interventions is still a subject of debate (Brunoni and Vanderhasselt 2014; Horvath et al. 2015; Hill et al. 2016; Mancuso et al. 2016; Wischnewski et al. 2021). The lack of efficient, scalable, and non-pharmacological interventions to improve WM highlights the need for developing new therapeutic strategies that directly target the underlying brain basis.

Real-time functional magnetic resonance imaging (rtfMRI) neurofeedback has been increasingly employed as a novel non-invasive brain modulation technique that capitalizes on the blood-oxygen-level dependent (BOLD) signal to allow individuals to gain volitional control over regional brain activity or connectivity and to modulate behavioral and cognitive functions associated with the respective brain systems (Zhao, Yao, et al. 2019; Zweerings et al. 2020; Goldway et al. 2022), including the enhancement of cognitive functions (Zhang et al. 2013; Scharnowski and Weiskopf 2015; Sherwood et al. 2016; Thibault et al. 2018; Da Silva and De Souza 2021). In contrast to pharmacological interventions, rtfMRI neurofeedback is comparably safe, has few negative side effects, and initial studies suggest that the effects can be maintained for several days after the training, even following a single training session (Hawkinson et al. 2012; Yao et al. 2016; Watanabe et al. 2017; Zhao, Yao, et al. 2019; Pamplona et al. 2020). Relatedly, functional near-infrared spectroscopy (fNIRS) is a (relatively) recently developed non-invasive optical brain imaging technique that captures changes in neural activity by detecting blood oxygenation levels by means of near-infrared light. The technique has been increasingly used in experimental settings, including real-time fNIRS (rtfNIRS) neurofeedback studies. Although rtfNIRS, compared to rtfMRI, has some limitations (e.g., restricted penetration depth and spatial resolution (Li et al. 2019; Kohl et al. 2020)), it also offers many practical advantages over fMRI, such as low cost, ability to measure neural activity in natural contexts, portability, and low sensitivity to motion artifacts. These advantages are essential to successfully translate and develop rtfMRI-neurofeedback training approaches into a scalable intervention (Li et al. 2019; Kohl et al. 2020; Soekadar et al. 2021).

WM is closely associated with well-characterized fronto-parietal brain networks (Linden 2007). Several studies reported that activation in the dorsolateral prefrontal cortex (DLPFC) is positively associated with WM load, and studies in patients with left DLPFC lesions additionally unveiled a critical role of this region in manipulating WM information (Edin et al. 2009; Klingberg 2010; Barbey et al. 2013; Balderston et al. 2020). Thus, the DLPFC is a crucial hub for WM, and, in particular, the left DLPFC is thought to support verbal WM (Curtis and D’Esposito 2003; Andrews et al. 2011; Barbey et al. 2013; Hudak et al. 2017; Nouchi et al. 2021; Wu et al. 2022). A growing body of (fMRI and fNIRS) neurofeedback training studies has targeted the DLPFC and demonstrated that participants can learn to control activity in this region following longer training protocols (> = 2 sessions) (Zhang et al. 2013; Hosseini et al. 2016; Sherwood et al. 2016; Hudak et al. 2017; Nouchi et al. 2021; Wu et al. 2022). Both neurofeedback training techniques showed that successful modulation of this region can enhance WM performance (Zhang et al. 2013; Hosseini et al. 2016; Sherwood et al. 2016; Nouchi et al. 2021; Wu et al. 2022), with the exception of one NIRS study that did not find any changes possible due to a ceiling effect (Hudak et al. 2017). Evidence suggests that a single session of neurofeedback training can also improve the regulation of prefrontal cortex (PFC) activation and cognitive abilities in healthy and overweight (and obese) subjects (Kohl et al. 2019; Li et al. 2019; Garrison et al. 2021). However, it is not yet clear if one session of rtfNIRS neurofeedback training targeting the DLPFC is effective for healthy subjects, nor whether the neural training success translates (i.e., transfers) to behavioral and neural effects during the subsequent engagement in a WM task in the absence of feedback. Both aspects are critical for the translation of rtfNIRS neurofeedback training into novel interventions that could impact behavioral and neural functioning in everyday life.

Previous studies have evaluated the behavioral and neural effects of behavioral WM training. For example, a meta-analysis on WM cognitive training revealed differences in neural activation for trained tasks versus untrained transfer tasks. During trained tasks, decreased activation in parietal and visual areas and increased PFC and striatal activation have been observed. In contrast, increased striatal and inferior frontal gyrus activation, and decreased DLPFC activation have been observed for (untrained) transfer tasks (Salmi et al. 2018). Moreover, both WM and neurofeedback training typically not only depend on the specific target region, but rather on a complex interplay of different brain networks, such as the central executive network, default mode network, and salience network (Haller et al. 2013; Zhang, Zhang, et al. 2015; Emmert et al. 2016; Salmi et al. 2018; Kohl et al. 2019). To understand the specific brain changes that mediate the translation of fNIRS neurofeedback success to (untrained) WM tasks, and to further determine how such translation depends on network-level changes, here we also collected fMRI data during (untrained) WM transfer task performance.

Against this background, the present study conducted an fNIRS-informed neurofeedback training of the left DLPFC (N = 62 healthy male individuals) using a randomized sham-controlled between-subject design, followed by a subsequent evaluation of the behavioral and neural transfer effects by means of a WM paradigm and concomitant fMRI data acquisition. Specific aims of the study were to (1) investigate if participants can learn to increase their DLPFC activity in a single session of rtfNIRS neurofeedback training, as well as to (2) evaluate the neurofeedback transfer effects on WM processing on the behavioral and neural (activity and network) level. We hypothesized that (1) healthy participants can increase DLPFC activity over the course of one session of DLPFC neurofeedback training, and (2) the training would lead to improved WM performance and DLPFC engagement.

## 2. Materials and Methods

### 2.1. Participants and Procedures

62 healthy male participants with no self-reported current or past psychiatric, neurological, or other medical conditions were included. To control for sex differences and reduce potential menstrual cycle effects, we only enrolled male participants in this study (similar to Li et al. 2019). Participants were asked to abstain from smoking, alcohol, caffeine, and tea for 24 hours prior to the experiment. On the day of the training, eligible participants underwent (1) the assessment of a range of potential confounders related to motivation, emotional processing, and cognitive performance, (2) random assignment to left DLPFC or sham (yoke feedback) rtfNIRS-guided neurofeedback training, (3) a WM task with fMRI. Assessment of potential confounders included: State Anxiety Inventory (STAI) (Spielberger 1970), Beck Depression Inventory-II (BDI-II) (Beck et al. 1996); and Behavioral Inhibition and Behavioral Activation System (measured using the BIS/BAS scales) (Carver and White 1994). Before and after training the Positive and Negative Affect Schedule (PANAS) (Watson et al. 1988) and the Digit Symbol Substitution Test (DSST) were administered to examine unspecific effects of training on basal attention and emotional states (Kaufman 1983). After the rtfNIRS training, all participants rated their level of regulatiomn success on a 0 - 9 Likert scale to evaluate blinding in the sham group.

Data from four participants were excluded due to technical problems during fNIRS (N = 2) or the fMRI assessments (N = 2) or abnormal WM performance during fMRI (> ±3 SD from the other participants; n=1) resulting in a final sample size of N = 58 (based on the randomized training assignment and following exclusion: neurofeedback, NF group: N = 31, and sham feedback, SH group: N = 27). Please note that one participant fulfilled two exclusion criteria, therefore, a total of four participants were excluded. All participants provided written informed consent and the project was approved by the local ethics committee at the University of Science and Technology of China and in line with the latest revision of the Declaration of Helsinki.

### 2.2. Neurofeedback training protocols and procedures

During the fNIRS-training experiment, participants were informed that the goal of the training was to learn how to increase brain activity in the target region. Following the fitting of the fNIRS cap and testing the online signal quality, participants were placed in front of the neurofeedback monitor. To reduce trial and error attempts during the initial training, participants completed a pretraining phase to explore the most appropriate strategies for regulating their brain activity before the formal experiment. After participants reported that they had developed effective strategies for increasing brain activity (4 - 8 blocks), and a short recovery break of around 3 - 5 minutes, the formal experiment started.

Following the pre-training period, participants were randomly assigned to receive either real-time neurofeedback from the lDLPFC (channel 6) or sham feedback (yoke feedback from a previous participant). During neurofeedback training, online neurofeedback was provided using a graphical interface that presented a stone laying on a beach. The level of activity in the target channel was coupled with the height of the stone. Participants were asked to try to “lift the stone” on the beach as high as possible using the strategies they developed during the pretraining phase and were informed that floating height of the stone indicated the level of neural activity in our specific feedback channel (channel 6). The training session included four subsequent runs of alternating rest and regulation blocks (four blocks per run, block duration = 25 s). Participants were also asked not to adjust their breathing or head position (or body) in an attempt to affect the feedback signal. Red or green lights on top of the display screen implied the task type: the red light indicated a “rest” condition, during which participants were instructed to relax, while the green light indicated a “regulation” condition, where participants were instructed to lift the stone by controlling their brain activity. For a schematic overview please see Figure 1A.

**Figure 1.**
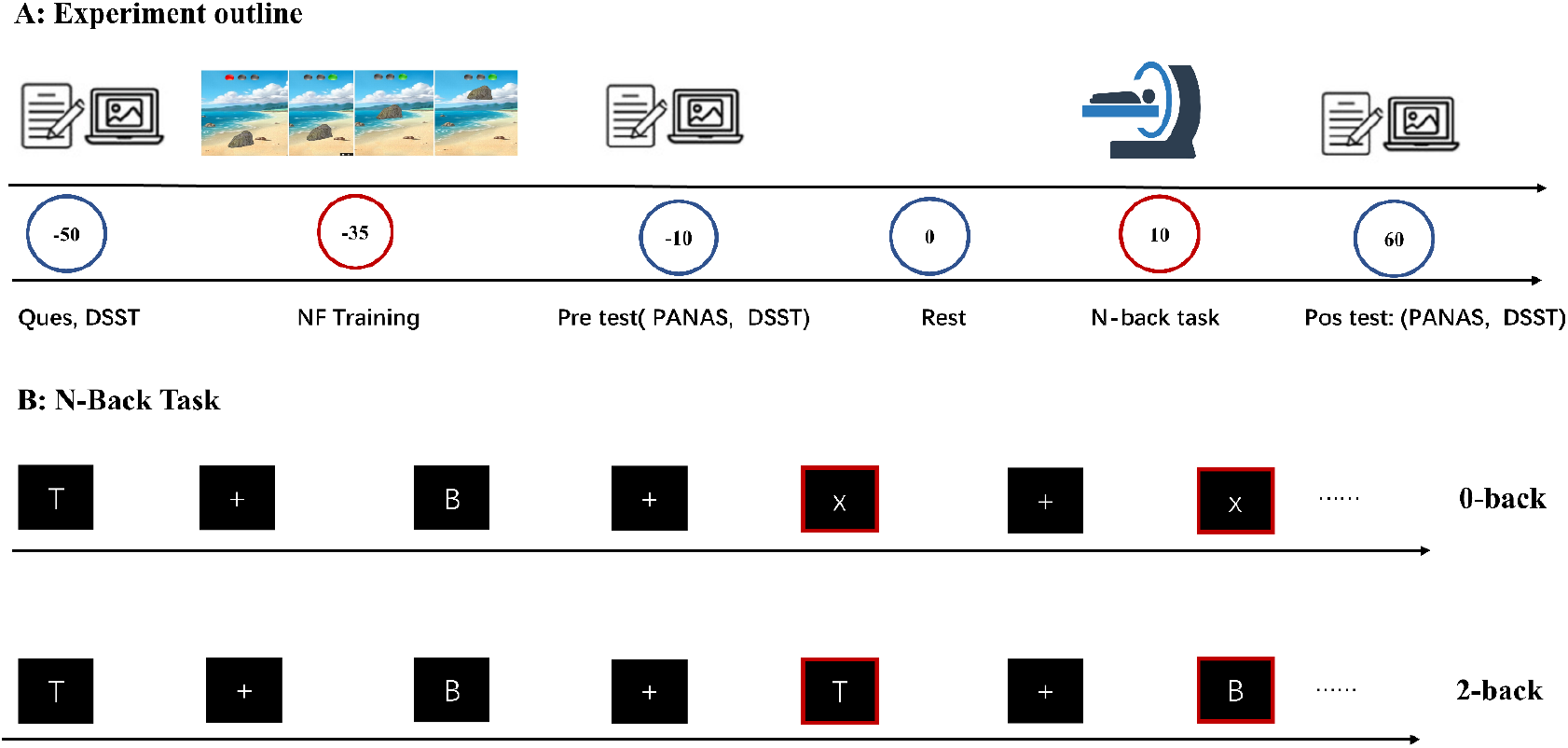
A, B: Experimental outline and schematic presentation of the N-back task including two levels of difficulty (0-back, 2-back).

### 2.3 NIRS data acquisition and real-time neurofeedback data preprocessing

Hemodynamic response signals were acquired using the NIRSport fNIRS system (NIRx Medizintechnik, GmbH, Berlin, Germany) with measures at two wavelengths (760 and 850nm) and a 7.81Hz sampling rate. The montage design used an optode set with 8 sources and 8 detectors, resulting in 18 source-detector pairs (channels). The optodes were placed according to the International 10-20 system for EEG electrode placement with the distance between the sources and detectors being 30 mm (Jasper 1958; Schommartz et al. 2020) (see Figure 2, note: channel 6 served as the target channel).

**Figure 2:**
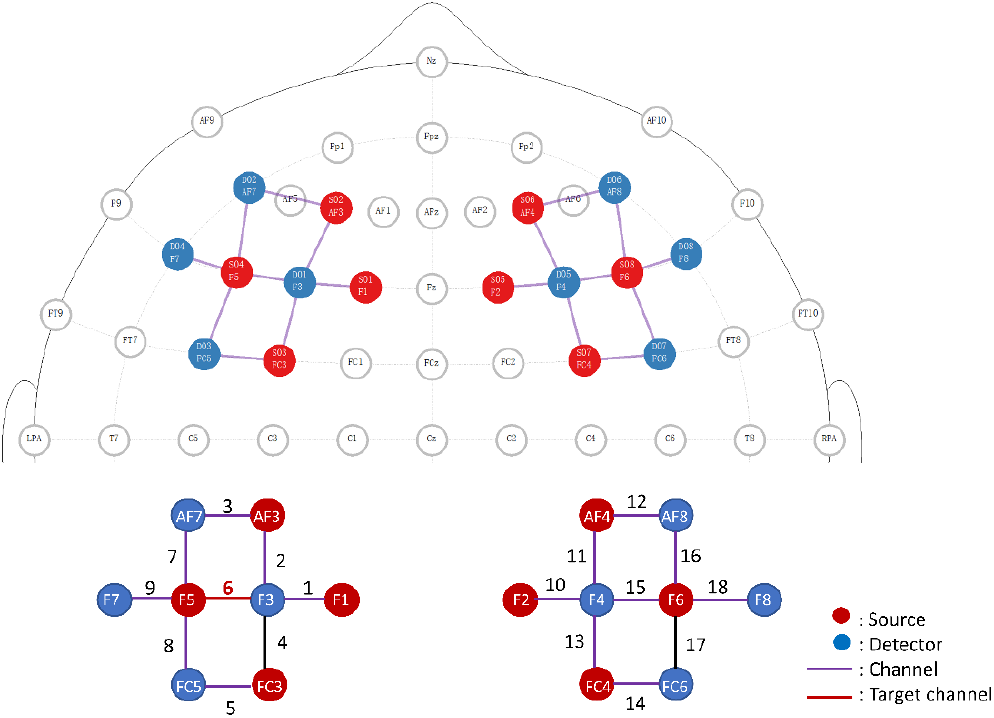
Montage of the dorsolateral prefrontal cortex: Source (S) and detectors (D) were positioned according to the international 10-20 system: Left hemisphere: Source: S1 at F1, S2 at AF3, S3 at FC3, S4 at F5, Detector: D1 at F3, D2 at AF7, D3 at FC5, D4 at F7; Right hemisphere: Source: S5 at F2, S6 at AF4, S7 at FC4, S8 at F6, Detector: D5 at F4, D6 at AF8, D7 at FC6, and D8 at F8.

The relative amplitude changes of oxygenated hemoglobin (Hbo) were used to calculate the feedback index. We used Hbo because it has a higher signal-to-noise ratio than deoxygenated hemoglobin (deoxy-Hbo) (Yamamoto and Kato 2002; Pinti et al. 2020). As a baseline measure of Hbo, we used the mean value of the 2 seconds before each regulation block and the raw Hbo signal was smoothed using a 2-second moving average window. So the realtime Hbo (f) was calculated by subtracting the preceding baseline Hbo from the smoothed Hbo signal and dividing it by the pre-defined difficulty coefficient: “M” (target channel 6 using the formula 1: 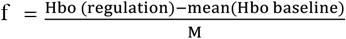). The difficulty coefficient was used to calibrate the feedback visualization based on the signal range of the current study design and target regions to improve the learning success of training (e,g., the small range of feedback visualization may not provide sufficient information to promote succesful neurofeedback learning). We determined the final feedback index (F), which ranges from 0 to 1 according to the range of the stone (bottom to top of the screen), using formula 1, which considers the relative change in brain activity 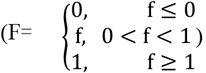. The calculated neurofeedback index (F) was displayed in a well-evaluated fNIRS neurofeedback platform (Li et al. 2019; Hou, Xiao, et al. 2021; Tang et al. 2021; Yu et al. 2021). Pilot assessments were conducted in independent experiments to examine the effective target channel (channel 6) and difficulty coefficient (M = 0.08; see section 2.3: Formula 1 and Figure 2 for detailed information, see also Li et al., (Li et al. 2019)), which was determined as optimal protocol parameter for voluntarily controlling activity in the left DLPFC.

### 2.4 WM task fMRI data acquisition

After the rtfNIRS neurofeedback training, participants completed three runs of a well-validated N-back WM task with concomitant fMRI. MRI data were collected on a 3.0 Tesla system (GE MR750, General Electric Medical system). Functional images were acquired using a T2*-weighted echo planar image sequence (repetition time [TR] = 2000 ms; echo time [TE] = 30 ms; flip angle [FA] = 90°; field of view [FOV]= 240 × 240 mm; resolution = 64 × 64; voxel size = 3.75 × 3.75 × 4 mm; slices number = 39; slice thickness = 1 mm).

A previously-validated N-back block-design paradigm with a 0-back and a 2-back condition was employed (Smith et al. 2018; Klugah-Brown et al. 2022). During the WM task, participants were asked to identify whether the current letter matched a specified target letter (X: 0-back condition) or the letter presented two before (2-back condition). We used phonologically closed letters to minimize the use of visual and phonological strategies (include stimuli: b, B, d, D, g, G, p, P, t, T, v, V). Each run included four blocks (2 blocks for 0-back condition and 2 blocks for 2-back condition). Each block started with the instruction presented for 2000 milliseconds to indicate the condition (0-back, 2-back), followed by a 4000 milliseconds blank screen before the first letter was presented. The letters were presented for 500 milliseconds with a 1500 millisecond fixed interval between them, blocks were separated by 8000 milliseconds of a fixation cross (see Figure 1 B).

### 2.5 Neural data analysis

#### 2.5.1 FNIRS offline data analysis

The NIRS-KIT was utilized for offline analysis of the neurofeedback data (Hou, Zhang, et al. 2021). Firstly the raw fNIRS signals were visually examined and corrected for motion artifacts with a temporal derivative distribution repair (TDDR) algorithm in case head movement was detected (Fishburn et al. 2019). We used a second-order polynomial regression model to decrease linear trends. Low-frequency and high-frequency physiological noise were reduced through the application of a band-pass filter (0.01-0.3 Hz) (Oda et al. 2018; Balconi and Fronda 2020). We additionally applied a systemized noise removal method to improve the reliability of fNIRS processing (Yamada et al. 2012). Finally, activity (beta value) in each channel was estimated using a general linear model (GLM) that included the two corresponding regressors (regulation and rest) and convolved regressors with standard hemodynamic response function (HRF) in each run (Friston et al. 1994).

#### 2.5.2 FMRI data analysis

The fMRI data were preprocessed and analyzed in SPM 12 (Statistical Parametric Mapping; http://www.fil.ion.ucl.ac.uk/spm/; Wellcome Trust Centre for Neuroimaging). Preprocessing steps included: 1) discarding the first five volumes to ensure MRI signal equilibrium, 2) slice time correction, 3) head motion correction, 4) normalization (resampled at 2 × 2 × 2 mm) using the EPI approach (Calhoun et al. 2017), and 5) spatial smoothing (8 mm full-width at half maximum [FWHM] Gaussian kernel). No participant was excluded due to head motion (max. head motion > 2.5 mm translation, > 2.5° rotation). Additionally, no significant differences in the average framewise displacement were found between groups (NF: 0.08 ±0.02; SH: 0.10 ±0.07, t (57) = 0.19) (Jenkinson et al. 2002).

To explore the effect of rtfNIRS neurofeedback training on WM neural activity, the two conditions (0 and 2-back), periods of task instructions, and fixation that separate each block were modeled separately as regressors and convolved with standard HRF. The six head motion parameters were included as covariates in the GLM. We initially examined the main effect of WM in the entire group (contrast: 2-back > 0-back) to verify the engagement of neural regions associated WM performance. Based on our a priori regional hypothesis and the training target region, we focused on the DLPFC to examine the effects of training on WM-related neural activity. The DLPF was therefore functionally defined using a meta-analytic approach in Neurosynth (https://neurosvnth.org/analvses/terms/dlpfc/). The corresponding peaks in the left and right DLPFC (left DLPFC: −38, 44, 10, right DLPFC: 38, 44, 10) were employed as the center of 15mm spheres to extract activity estimates. These were subjected to a mixed ANOVA with the neurofeedback training group (NF, SH) as a between-subject factor and condition (0-back, 2-back) as a within-subject factor.

#### 2.5.3 Training effect on functional connectivity of the target region

We further performed an exploratory generalized form of context-dependent psychophysiological interaction analysis (gPPI) to explore whether neurofeedback training could change the connectivity of the left DLPF on the network level (McLaren et al. 2012). The left DLPFC peak coordinate (10 mm sphere) identified in the group level analysis (peak MNI of lDLPFC: −48 36 8) was used as a seed region in a seed-to-whole brain functional connectivity analysis. The model included identical regressors as the BOLD activation design matrix as well as the corresponding psychophysiological interaction and the six motion regressors of no interest. We modeled context-dependent variations in functional connectivity from left DLPFC to the whole brain and examined voxel-level group differences (i.e., training-induced changes) using an exploratory two-sample t-test (contrast: 2-back > 0-back) with a threshold of p < 0.001 (uncorrected) and a cluster extent of > 50 voxels.

## 3. Results

### 3.1 Demographics and potential confounders

The NF and SH groups did not significantly differ in demographic data and potential confounders, with the exception of a marginal group difference in the average depression load as measured by the BDI-II (NF: 4.61 ± 3.69; SH: 7.93 ± 7.87, p = 0.04). BDI-II scores were consequently included as a covariate in all subsequent analyses. The NF and SH groups also did not exhibit differences in attentional performance (i.e., DSST performance) and positive/negative affect before or after the training (ps >0.05), suggesting an absence of training-induced unspecific effects on basal attention or mood. No group differences were observed with respect to the self-perceived training success (ps > 0.05) confirming successful blinding in the SH group.

### 3.2 fNIRS-informed training of lDLPFC activity – single session effects

A mixed two-way ANOVA was conducted to examine the effect of the training runs (run1, run2, run3, run4) and training (NF, SH) on Hbo values from each channel. We observed a main effect of run in the lDLPFC target channel (channel 6: F(3,165) = 3.72, p = 0.01, η^2^ = 0.04), as well as a significant interaction between run and group in the target channel and its adjacent channels (channel 5: F(3,165) = 3.46, p = 0.02, η^2^ = 0.05; channel 6: F(3,165) = 3.20, p = 0.03, η^2^ = 0.03; channel 7: (3,165) = 2.71, p = 0.05, η^2^ = 0.04; channel 8: F(3,165) = 2.84, p = 0.04, η^2^ =0.04), suggesting that the NF and SH group exhibited different Hbo values as the experiment progressed. No significant main or interaction effects were observed for the other channels (all ps > 0.05).

One-way ANCOVAs were applied to further test training-induced changes over the runs (run1, run2, run3, run4) within each training group. In these analyses, we observed significant main effects of run in the NF group in training channel 6, and adjacent channels 4 and 5 (channel 4: F(3,119) = 2.88, p = 0.04, η^2^ = 0.07, channel 5: F(3,119) = 3.81, p = 0.01, η^2^ = 0.09, channel 6: F (3,119) = 4.03, p = 0.009, η^2^ = 0.09). Post-hoc tests further showed that the Hbo signal in the lDLPFC significantly increased over the runs in the NF group for both the target and adjacent channels (channel 2: run 1 VS run 4: p_FDR_ = 0.01; channel 3: run 1 VS run 4: p_FDR_ = 0.01; channel 4, run1 VS run 4: p_FDR_ = 0.01; channel 5, run1 VS run 4: p _FDR_ = 0.09; channel 6, run1 VS run 4: p _FDR_ = 0.001; FDR correction was employed accounting for the number of all channels to account for multiple comparions). Direct comparisons of the NF and SH groups for the individual runs, however, did not reveal significant between-group differences in any channel (P _FDR_ > 0.05). Together these results suggest that the NF training induced an increase in Hbo from run 1 to 4 in the target and adjacent channels.

In the SH group, a significant main effect of run was found in channel 10 (F(3,104) = 2.93, p = 0.04, η^2^ = 0.08), a channel located contra-lateral to the training target channel over the right DLPFC and the adjacent right ventrolateral prefrontal cortex. Post-hoc tests further confirmed an increase in channel 10 over the training runs in the SH group, but results were not significant after FDR correction (run 2 VS run 4: p_uncorrected_ = 0.04) (Figure 3).

**Figure 3.**
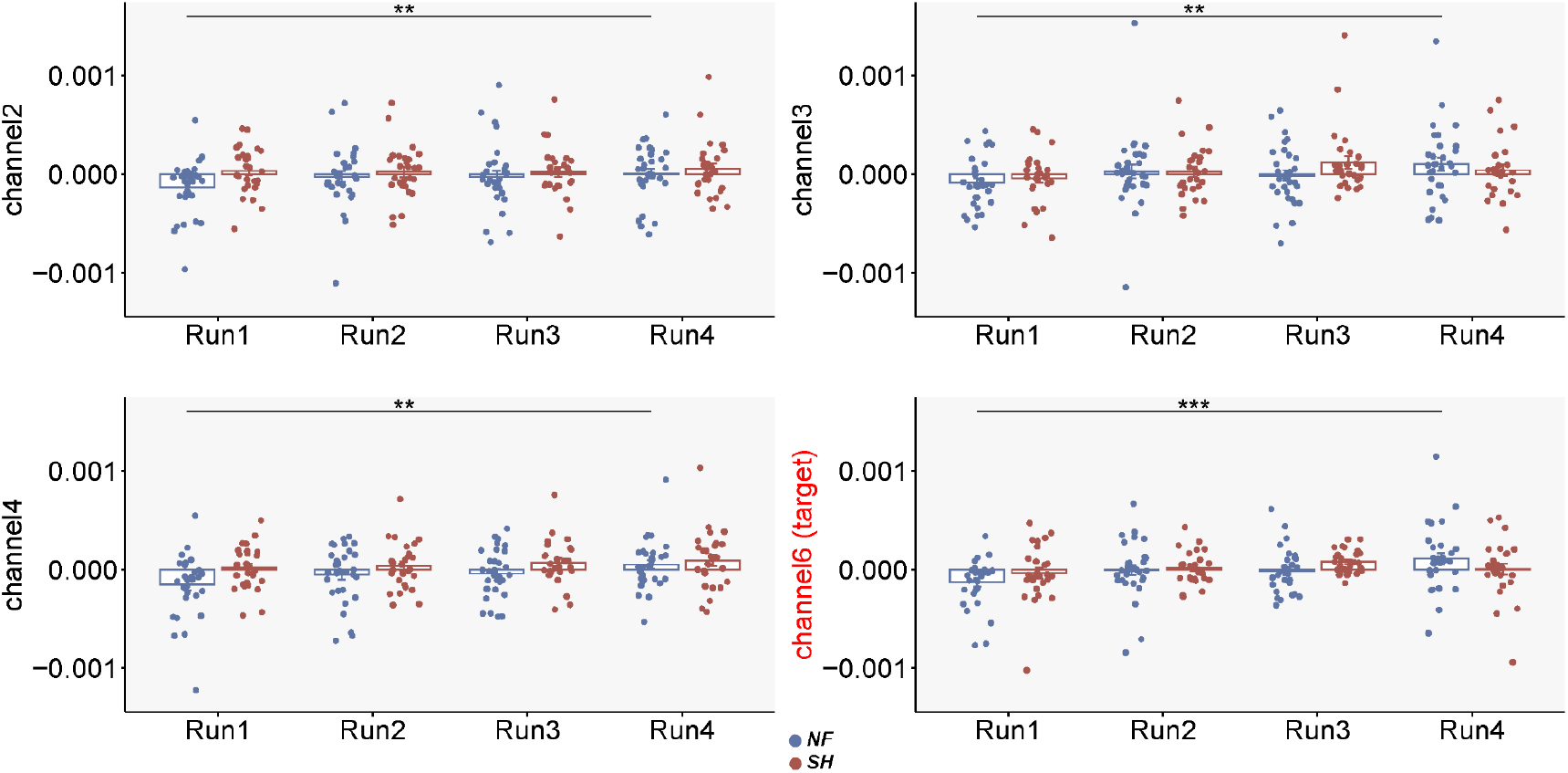
Post hoc tests (run 1 VS run 4) indicated that the Hbo signal in channels 2, 3,4 and channel 6 (target channel) significantly increased over the training runs in the NF group. NF: neurofeedback group, SH: sham feedback group. **, p_FDR_ ≤ 0.01, ***, p_FDR_ ≤ 0.001.

### 3.3 Transfer effects of training on the behavioral level – WM performance

In mixed ANOVA analyses with the training groups (NF, SH) as a between-subject factor and N-back conditions (0-back, 2-back) as within-subject factor, we did not observe a significant main or interaction effect on reaction times (RT) or accuracy (ACC) (all ps > 0.15). However, a significant main effect of condition (ACC: F (1,55) = 16.95, p < 0.001, η^2^ = 0.03; RT (1,55) = 60.57, p < 0.001, η^2^=0.03) was observed for both outcome measures. Specifically, participants performed better (faster RT and higher ACC) under the 0-back load level compared to the 2-back load level (ACC: p_Bonferroni corrected_ < 0.001, Cohen’s d 0.57; RT. p_Bonferroni corrected_ < 0.001, Cohen’s d = 0.45) reflecting a successful WM load manipulation.

### 3.4 Evidence for the engagement of WM-related brain networks during N-back

We initially replicated previous findings suggesting that the N-back task induces robust activation increases in frontoparietal processing networks in the entire group (contrast: 2-back > 0-back), including the medial frontal gyrus, left inferior frontal gyrus/insula and left inferior frontal gyrus/precentral, left parietal lobe, left precuneus, right inferior parietal lobule left precuneus, right insula, right cerebellum, right inferior frontal gyrus, right middle frontal gyrus (Wager and Smith 2003; Emch et al. 2019) (see Supplementary Figure 1 and Table 1).

**Table 1:**
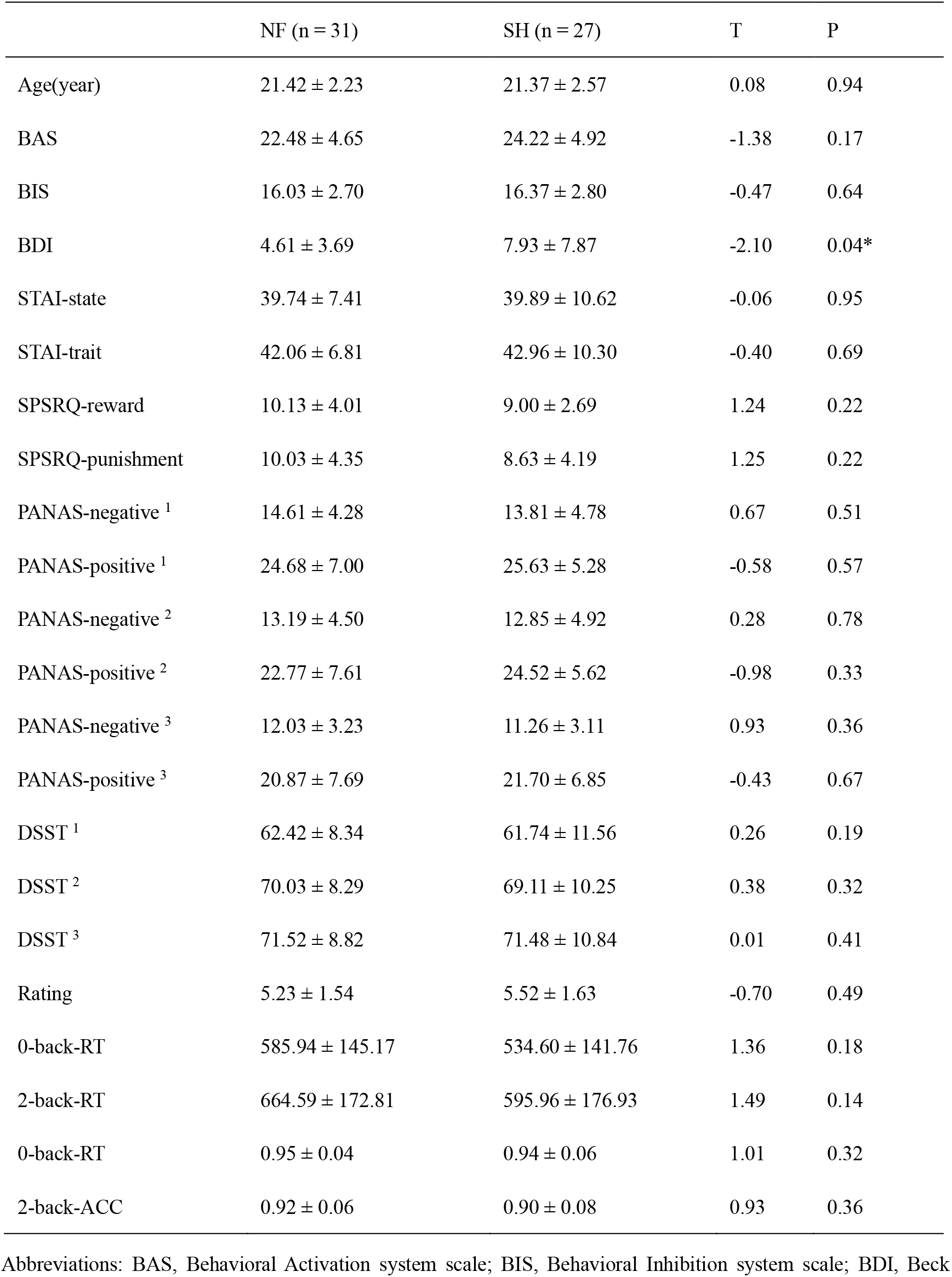
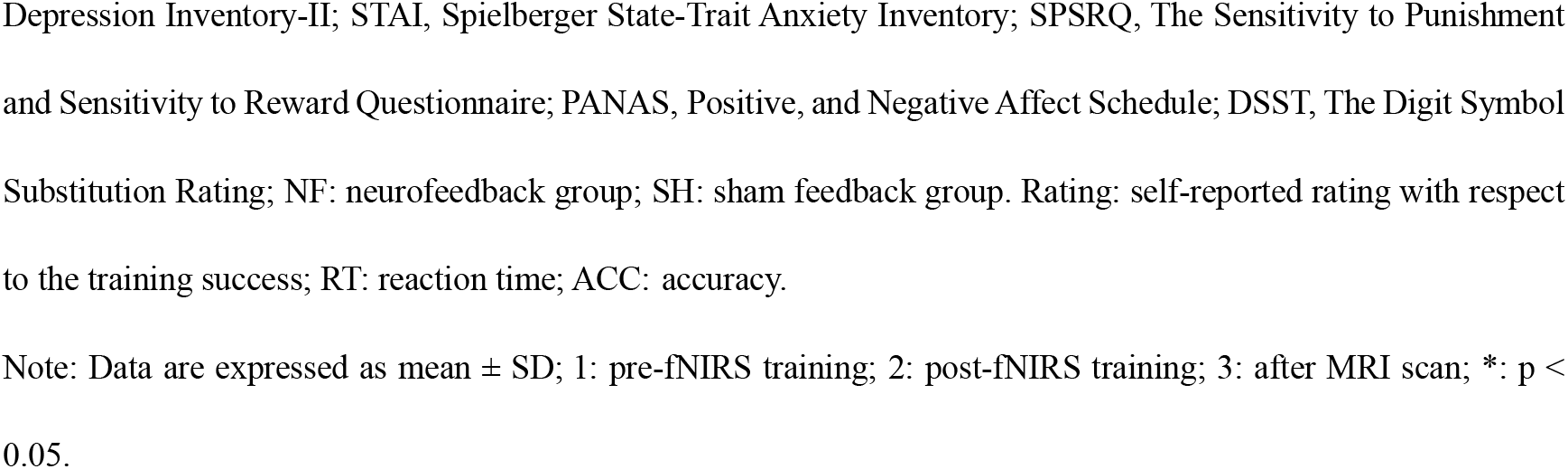
Descriptive statistics of the experimental groups.

|  | NF (n = 31) | SH (n = 27) | T | P |
| --- | --- | --- | --- | --- |
| Age(year) | 21.42 ± 2.23 | 21.37 ± 2.57 | 0.08 | 0.94 |
| BAS | 22.48 ± 4.65 | 24.22 ± 4.92 | -1.38 | 0.17 |
| BIS | 16.03 ± 2.70 | 16.37 ± 2.80 | -0.47 | 0.64 |
| BDI | 4.61 ± 3.69 | 7.93 ± 7.87 | -2.10 | 0.04* |
| STAI-state | 39.74 ± 7.41 | 39.89 ± 10.62 | -0.06 | 0.95 |
| STAI-trait | 42.06 ± 6.81 | 42.96 ± 10.30 | -0.40 | 0.69 |
| SPSRQ-reward | 10.13 ± 4.01 | 9.00 ± 2.69 | 1.24 | 0.22 |
| SPSRQ-punishment | 10.03 ± 4.35 | 8.63 ± 4.19 | 1.25 | 0.22 |
| PANAS-negative <sup>1</sup> | 14.61 ± 4.28 | 13.81 ± 4.78 | 0.67 | 0.51 |
| PANAS-positive <sup>1</sup> | 24.68 ± 7.00 | 25.63 ± 5.28 | -0.58 | 0.57 |
| PANAS-negative <sup>2</sup> | 13.19 ± 4.50 | 12.85 ± 4.92 | 0.28 | 0.78 |
| PANAS-positive <sup>2</sup> | 22.77 ± 7.61 | 24.52 ± 5.62 | -0.98 | 0.33 |
| PANAS-negative <sup>3</sup> | 12.03 ± 3.23 | 11.26 ± 3.11 | 0.93 | 0.36 |
| PANAS-positive <sup>3</sup> | 20.87 ± 7.69 | 21.70 ± 6.85 | -0.43 | 0.67 |
| DSST <sup>1</sup> | 62.42 ± 8.34 | 61.74 ± 11.56 | 0.26 | 0.19 |
| DSST <sup>2</sup> | 70.03 ± 8.29 | 69.11 ± 10.25 | 0.38 | 0.32 |
| DSST <sup>3</sup> | 71.52 ± 8.82 | 71.48 ± 10.84 | 0.01 | 0.41 |
| Rating | 5.23 ± 1.54 | 5.52 ± 1.63 | -0.70 | 0.49 |
| 0-back-RT | 585.94 ± 145.17 | 534.60 ± 141.76 | 1.36 | 0.18 |
| 2-back-RT | 664.59 ± 172.81 | 595.96 ± 176.93 | 1.49 | 0.14 |
| 0-back-RT | 0.95 ± 0.04 | 0.94 ± 0.06 | 1.01 | 0.32 |
| 2-back-ACC | 0.92 ± 0.06 | 0.90 ± 0.08 | 0.93 | 0.36 |

Abbreviations: BAS, Behavioral Activation system scale; BIS, Behavioral Inhibition system scale; BDI, Beck
Depression Inventory-II; STAI, Spielberger State-Trait Anxiety Inventory; SPSRQ, The Sensitivity to Punishment and Sensitivity to Reward Questionnaire; PANAS, Positive, and Negative Affect Schedule; DSST, The Digit Symbol Substitution Rating; NF: neurofeedback group; SH: sham feedback group. Rating: self-reported rating with respect to the training success; RT: reaction time; ACC: accuracy.
Note: Data are expressed as mean ± SD; 1: pre-fNIRS training; 2: post-fNIRS training; 3: after MRI scan; \*: $p < 0.05$ .

### 3.5 Transfer effects of training on regional brain activity – WM-related DLPFC activation

Examination of neural activity in the left and right DLPFC using a mixed ANOVA with group and WM load revealed a significant main effect of condition and group in the left DLPFC (condition: F(1,56) = 35.30, p < 0.001, η^2^ = 0.12; group: F(1,56) = 6.64, p = 0.01, η^2^ = 0.07), and, for the right DLPFC, a significant main effect of group (F(1,56) = 5.38, p = 0.02, η^2^ = 0.07) and significant group-by-condition interaction (F(1,56) = 3.88, p = 0.05, η^2^ = 0.01), with the interaction effect suggesting that compared to sham feedback, neurofeedback training induced a greater increase between 0-back and the 2-back load level. Post hoc tests indicated that the NF group exhibited higher bilateral DLPFC activity compared to the SH group across both WM load conditions, but comparisons only survived in 2-back load level after multiple comparison corrections (0-back-1DLPFC: p_not corrected_ = 0.03, Cohen’s d = 0.51; 0-back-rDLPFC: p_not corrected_ = 0.21, Cohen’s d = 0.31; 2-back-1DLPFC: p_Bonferroni corrected_ = 0.07, Cohen’s d = 0.67; 2-back-rDLPFC, p_Bonferroni corrected_ = 0.02, Cohen’s d = 0.78) (see Figure 4 A).

**Figure 4.**
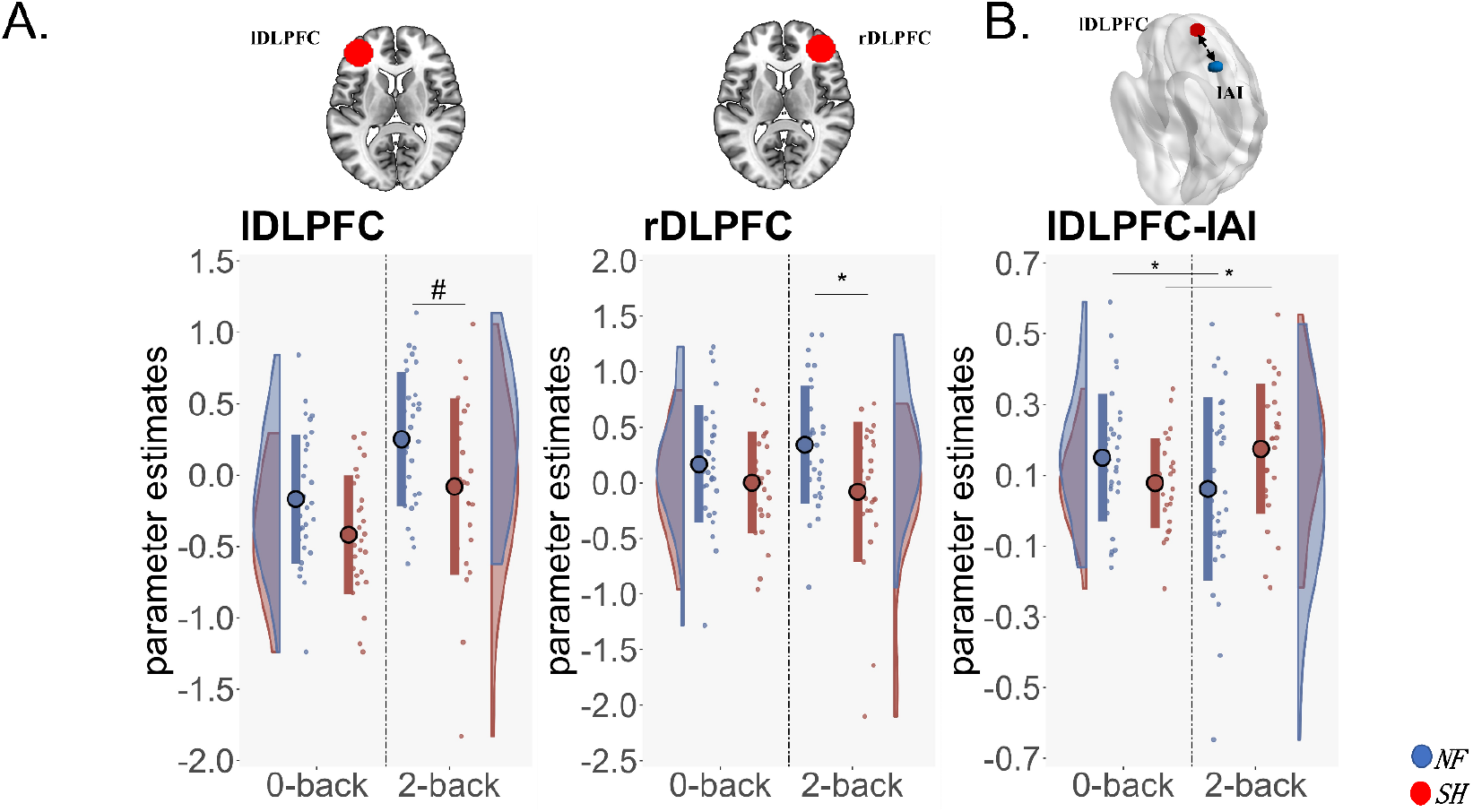
Neurofeedback training effects of rtfNIRS on WM-related DLPFC activation. (A) The left and right DLPFC showed increased activity during WM tasks (2-back load levels) in the NF training group as compared to the SH group. (B) NF training significantly decreased left DLPFC connectivity strength with the left AI, compared to SH during the WM task. NF: neurofeedback group, SH: sham feedback group, ldlPFC: the left dorsolateral prefrontal cortex, rdlPFC: the right dorsolateral prefrontal cortex, AI: anterior insula, WM: working memory task. #:p_Bonferroni corrected_ =0.07, *: p_Bonferroni corrected_ <0.05.

### 3.6 Transfer effects of training on the network level – WM-related DLPFC connectivity

The exploratory functional connectivity analysis revealed that, during WM processing (2-back > 0-back contrast), the NF group, relative to the SH group, exhibited decreased left DLPFC coupling with a cluster located in the left anterior insular (AI) cortex (peak MNI: −42 20 4, p uncorrected < 0.001, k = 68) (see Figure 4 B).

### 3.7 Associations between DLPFC activity and WM performance

We examined associations between the extracted DLPFC activity (from the ROIs used for evaluating the training effects on the fMRI data) and WM performance in each group separately. We observed a significant negative correlation between RT and bilateral DLPFC activity in the NF group (0-back-1DLPFC, r = −0.39, p = 0.03; 0-back-rDLPFC, r = −0.41, p = 0.02; 2-back-lDLPFC: r = −0.46, p = 0.009; 2-back-rDLPFC = −0.38, p = 0.04), but no significant correlations were found in the SH group. The difference between correlation coefficients was only marginally significant in the 2-back condition and corresponding DLPFC activity (z_2-back-lDLPFC = −1.79, one-tailed, p = 0.03, two-tailed, p=0.07; z 2-back-rDLPFC = −1.69, one-tailed, p = 0.05, two -tailed, p=0.09). These results may reflect that WM processing speed was significantly associated with DLPFC activity after rtfNIRS training (shown in Figure 5).

**Figure 5.**
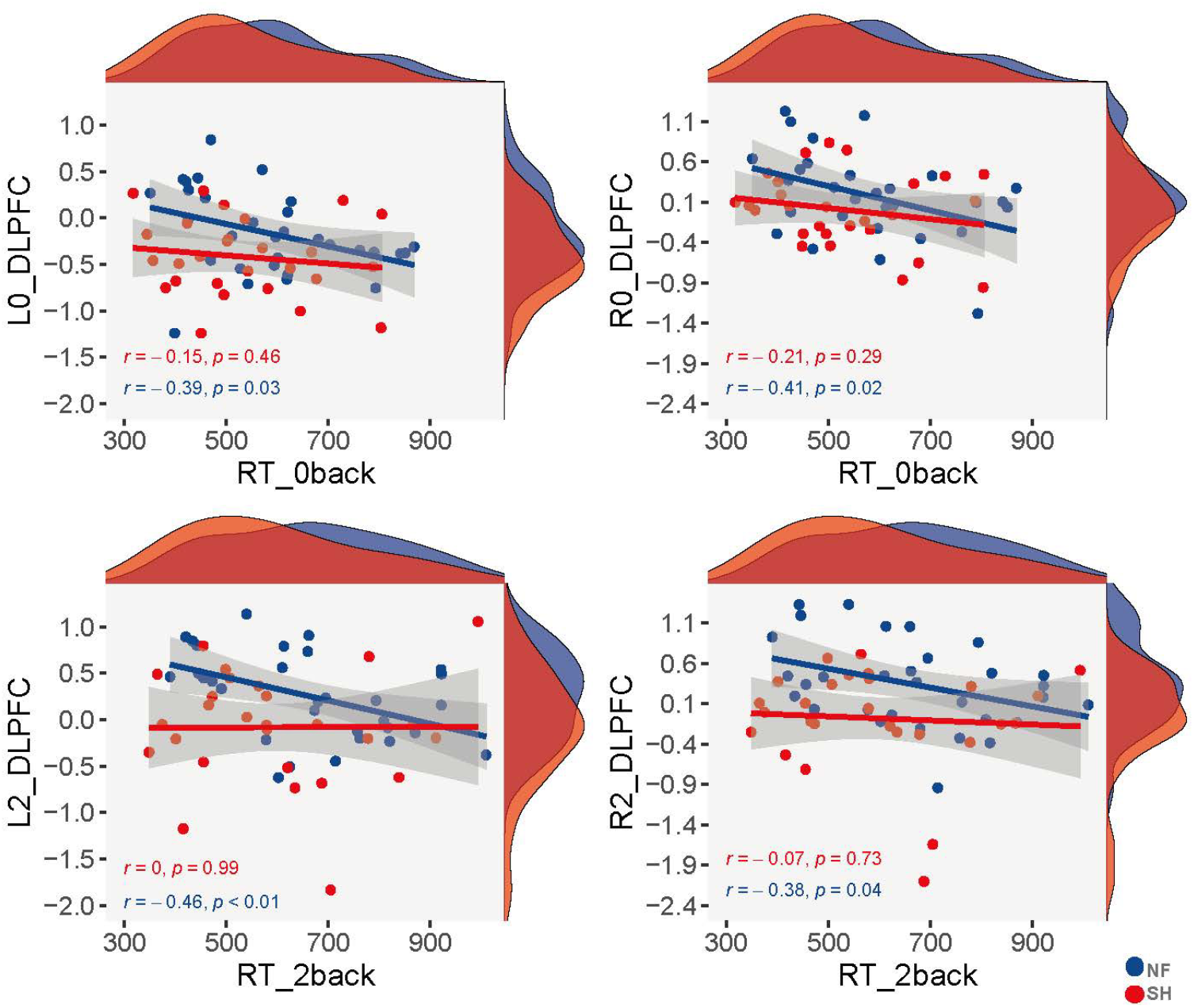
Associations between the DLPFC activity and RT in the working memory task: In the NF group, RT was significantly negatively correlated with bilateral DLPFC activity (all P < 0.05), whereas no significant association was observed in the SH group. The filled curves show standard errors for each group and the distribution curves in marginals indicate statistical densities of RT and bilateral DLPFC activity respectively. RT: reaction time. NF: neurofeedback group; SH: sham feedback group. The grey shaded area identifies the 95 % confidence interval.

## 4. Discussion

Here, we aimed to investigated whether a single session of neurofeedback training can induce behavioral and neural changes during WM processing in the absence of online feedback, using a randomized sham-controlled between-subjects study combining fNIRS-informed neurofeedback and an fMRI WM paradigm. Investigating such transfer effects is critical for the successful translation of lab-based NF studies to potentially therapeutic real-life interventions. In line with previous studies employing rtfNIRS neurofeedback to modulate lateral frontal brain activity (Hosseini et al. 2016; Li et al. 2019; Nouchi et al. 2021; Yu et al. 2021; Wu et al. 2022), participants in the present study were able to gain control over left DLPFC activation within a single (four-run) training session. Specifically, participants who were provided with rtfNIRS neurofeedback were able to successfully increase activity in the left DLPFC over the four training runs, while no corresponding changes in the same region were observed in the SH group. Neural changes in the neurofeedback group were mainly restricted to the target (i.e., DLPFC) and adjacent channels (i.e., left lateral PFC), suggesting that our training successfully induced regionally-specific effects. During the subsequent transfer assessment, in which we combined a well-validated N-back WM task with fMRI, we observed that the active and sham NF groups did not differ on any task performance parameters, including ACC and RTs. Importantly, however, the neurofeedback training group exhibited generally increased bilateral DLPFC activation during the task, as well as reduced WM-load dependent functional connectivity between left DLPFC and AI. Exploratory analyses further revealed that, within the active NF group, faster RTs during the WM task were associated with stronger training-induced DLPFC activity. Together, the present findings suggest that a single session of rtfNIRS neurofeedback training can enable participants to learn to volitionally up-regulate dlPFC activity and that such training effects are maintained during subsequent engagement in a cognitive task that critically depends on DLPFC function.

A number of previous neurofeedback studies demonstrated that healthy participants can learn to control activity in the DLPFC via online neurofeedback although most previous studies employed multiple sessions of training (Zhang et al. 2013; Marx et al. 2014; Hosseini et al. 2016; Sherwood et al. 2016; Hudak et al. 2017; Kimmig et al. 2019; Nouchi et al. 2021; Wu et al. 2022). The present study further underscores the potential of rtfNIRS neurofeedback as a novel approach to train healthy individuals to gain regulatory control over prefrontal brain activity and suggests that - at least healthy subjects - can rapidly learn to self-regulate DLPFC activation within a single training session. Given its low cost, user-friendly implementation, and ability to be implemented as a mobile system in everyday settings, rtfNIRS neurofeedback can facilitate the application of Hbo-activity-informed neurofeedback into therapeutic and practical applications (Kohl et al. 2020). For the translation into application, it is however critical that the behavioral and neural effects of the training are maintained in other contexts and in the absence of neurofeedback.

To evaluate the neural transfer effects in the absence of neurofeedback we employed fMRI. FNIRS technical restrictions do not allow measurements of brain activity below the cortical surface (Liu et al. 2015), which thus necessitates the use of fMRI to investigate possible transfer effects in deeper (sub-)cortical brain regions with higher spatial resolution (Emmert et al. 2016; Sitaram et al. 2017). In line with our hypothesis, the training group exhibited higher DLPFC activity during the WM task as compared to the SH group. These findings demonstrate that the training effects on the neural level were transferred to subsequent cognitive tasks and that rtfNIRS neurofeedback may have affected neural efficiency during cognitive processes (Hosseini et al. 2016; Sherwood et al. 2016; Hudak et al. 2017). Accumulating evidence shows that short-term WM training decreases DLPFC activity (Landau et al. 2004; Sayala et al. 2005; Landau et al. 2007; Brooks et al. 2020), while longer training increases DLPFC activity (Olesen et al. 2004; Westerberg and Klingberg 2007; Jolles et al. 2010; Klingberg 2010; McKendrick et al. 2014). Neuroenhancement studies using pharmacological agents as well as brain stimulation approaches commonly reported enhanced executive functioning in the context of enhanced prefrontal activity (Rasetti et al. 2010; Li et al. 2020) and previous rtfMRI-informed neurofeedback studies targeting the DLPFC found improved WM performance and enhanced DLPFC activity (Zhang et al. 2013; Shen et al. 2015; Zhang, Zhang, et al. 2015; Sherwood et al. 2016). Within this context, the present results may reflect that rtfNIRS-informed neurofeedback training success can translate into new cognitively demanding situations in the absence of online feedback. While we did not observe an effect on the behavioral level (although see our discussion below), the enhanced activation may have the potential to support cognitive functioning. The findings from the exploratory correlation analyses yielding a significant negative correlation between RTs and DLPFC may provide further tentative support for the behavioral potential of the training (Sligte et al. 2011; Hudak et al. 2017; Kumar et al. 2017; Nouchi et al. 2021).

On the network level, we observed - in an exploratory whole-brain analysis - that the NF group exhibited a load-dependent decrease in functional connectivity between the target region of the left DLPFC and the ipsilateral AI while the SH group demonstrated a load-dependent increase. Some previous studies reported network level changes following training of the DLPFC with fMRI - and one study with rtfNIRS neurofeedback training. These studies reported that training enhanced functional connectivity between the salience network and central executive and default mode regions (Shen et al. 2015; Zhang, Yao, et al. 2015; Zhang, Zhang, et al. 2015; Kohl et al. 2019; Yu et al. 2021), as well as decreased connectivity with the insula (Kohl et al. 2019). While differences in the training session, instruction and the specific target area impede a direct comparison of the findings, together the results underscore the potential of neurofeedback training that targets isolated activity in single regions to change also the communication on the network level and thus influence regions that are not directly trained. As a core region of the salience network, the AI has been consistently reported as a critical node that supports switching between external and internal task demands and a ‘gatekeeper to executive control’, it also plays a critical role in the reorganization of brain networks and improvements of WM after neurofeedback training (Zhang, Zhang, et al. 2015; Cai et al. 2021; Molnar-Szakacs and Uddin 2022). Previous studies reported that a stronger dissociation between the insular salience nodes and frontal-executive networks in an attention task was associated with better cognitive performance (Elton and Gao 2014), and thus the load-dependent decrease may reflect improved network level communication during WM engagement.

The absence of behavioral (i.e., task performance) transfer effects of NF training may suggest that more intensive and longer training is necessary to modulate cognitive functions such as WM (Weiss et al. 2022). Previous work has demonstrated that neurofeedback training effects begin to appear after two training sessions, but that they may require up to 30 sessions for rtfNIRS neurofeedback to reach their plateau (Zhang et al. 2013; Sherwood et al. 2016; Hudak et al. 2018; Rance et al. 2018; Kohl et al. 2020), indicating a clear need for longer and more intense training protocols (Rance et al. 2018). Moreover, WM is not a unitary function, as it incorporates a number of subprocesses including short-term storage and manipulation of information, both of which have been linked to DLPFC function (Daniel et al. 2016; Wang et al. 2019). While the present N-back task reliably engaged the WM networks, the inclusion of a single load condition (2-back) and the inherent focus of the task on the storage and updating component of WM, may have limited the task’s sensitivity to behavioral transfer effects.

This study also has some limitations. Firstly, our study only included male participants to reduce variance related to sex differences and future studies need to determine the generalizability of our results. Secondly, we did not observe behavioral transfer effects following the short-time rtfNIRS training. Future studies need to test whether more intensive training periods can induce behavioral improvements or whether effects in specific subdomains of WM can be observed. Thirdly, contamination of the fNIRS signal with superficial hemodynamic fluctuations reduces the signal-to-noise ratios of fNIRS data (Brigadoi and Cooper 2015), and new technologies such as short separation channels could be used in future studies to control for artifacts and may help to improve the signal-to-noise-ratio in further studies (Kohl et al. 2020).

## 5. Conclusion

Overall, the present study suggests that healthy participants can rapidly acquire control over the DLPFC activation via a single training of rtfNIRS-informed neurofeedback. Moreover, we demonstrate that such training effects can transfer to a subsequent working memory task, as evidenced by increased DLPFC activation during task engagement, and changes in functional coupling strength between left DLPFC and AI. While no behavioral transfer effects were observed, the presence of a negative association between training-induced DLPFC changes and faster RT in the active training group provides some preliminary evidence for the potential for inducing transfer effects on the behavioral level. More robust transfer effects may be acquired via using longer training protocols or more sensitive WM tasks. Overall, the results further underscore the application potential of rtfNIRS-neurofeedback. Given that WM impairments represent a transdiagnostic feature of several neurologic and psychiatric disorders, rtfNIRS training of DLPFC may represent a promising therapeutic strategy. Future work may therefore focus on exploring such effects in clinical populations.

## Author contributions

Conceptualization, Methodology: Xi Yang, and Benjamin Becker.

Performed the experiments: Xi Yang, Yixu Zeng, Guojuan Jiao.

Data analysis: Xi Yang, Xianyang Gan, Yihan Jiang.

Writing - original draft: Xi Yang and Benjamin Becker.

Writing - review & editing: Xi Yang, Dennis Hernaus, David Linden, Shuxia Yao, Dezhong Yao, and Benjamin Becker.

Technical support: Chaoze Zhu, Yihan Jiang.

Funding acquisition: Benjamin Becker.

## Ethics approval statement

The research was approved by the local ethics committee of the University of Electronic Science and Technology of China and all procedures were in compliance with the latest revision of the Declaration of Helsinki.

## Statement of human rights

Participants provided informed written consent after they were informed clearly of the detailed procedures.

## Acknowledgments

The authors would like to thank co-authors valuable help and we also want to appreciate all the participants in the present study.

## Data/code availability statement

The processed behavioral, fNIRS, and fMRI data and maps are available at the OSF project site (https://osf.io/hf4cr/).

## References

Afsaneh E, Zarei Ghobadi M. 2022. The role of neuromodulation-related technologies in neurology for the next 10 years. Brain-Apparatus Communication: A Journal of Bacomics.1–6.

Andrews SC, Hoy KE, Enticott PG, Daskalakis ZJ, Fitzgerald PB. 2011. Improving working memory: the effect of combining cognitive activity and anodal transcranial direct current stimulation to the left dorsolateral prefrontal cortex. Brain Stimul. 4:84–89.

Balconi M, Fronda G. 2020. Morality and management: an oxymoron? fNIRS and neuromanagement perspective explain us why things are not like this. Cognitive, Affective, & Behavioral Neuroscience. 20:1336–1348.

Balderston NL, Flook E, Hsiung A, Liu J, Thongarong A, Stahl S, Makhoul W, Sheline Y, Ernst M, Grillon C. 2020. Patients with anxiety disorders rely on bilateral dlPFC activation during verbal working memory. Soc Cogn Affect Neurosci. 15:1288–1298.

Barbey AK, Koenigs M, Grafman J. 2013. Dorsolateral prefrontal contributions to human working memory. cortex. 49:1195–1205.

Beck AT, Steer RA, Brown GK. 1996. Manual for the beck depression inventory-II. San Antonio, TX: Psychological Corporation. 1:10.1037.

Becker B, Wagner D, Gouzoulis-Mayfrank E, Spuentrup E, Daumann J. 2010. The impact of early-onset cannabis use on functional brain correlates of working memory. Prog Neuropsychopharmacol Biol Psychiatry. 34:837–845.

Brigadoi S, Cooper RJ. 2015. How short is short? Optimum source-detector distance for short-separation channels in functional near-infrared spectroscopy. Neurophotonics. 2:025005.

Brooks SJ, Mackenzie-Phelan R, Tully J, Schiöth HB. 2020. Review of the Neural Processes of Working Memory Training: Controlling the Impulse to Throw the Baby Out With the Bathwater. Front Psychiatry. 11:512761.

Brunoni AR, Vanderhasselt M-A. 2014. Working memory improvement with non-invasive brain stimulation of the dorsolateral prefrontal cortex: A systematic review and meta-analysis. Brain and Cognition. 86:1–9.

Cai W, Ryali S, Pasumarthy R, Talasila V, Menon V. 2021. Dynamic causal brain circuits during working memory and their functional controllability. Nature Communications. 12:3314.

Calhoun VD, Wager TD, Krishnan A, Rosch KS, Seymour KE, Nebel MB, Mostofsky SH, Nyalakanai P, Kiehl K. 2017. The impact of T1 versus EPI spatial normalization templates for fMRI data analyses. Hum Brain Mapp. 38:5331–5342.

Carver CS, White TL. 1994. Behavioral inhibition, behavioral activation, and affective responses to impending reward and punishment: the BIS/BAS scales. Journal of personality and social psychology. 67:319.

Curtis CE, D’Esposito M. 2003. Persistent activity in the prefrontal cortex during working memory. Trends in Cognitive Sciences. 7:415–423.

Da Silva JC, De Souza ML. 2021. Neurofeedback training for cognitive performance improvement in healthy subjects: A systematic review. Psychology & Neuroscience. 14:262.

Daniel TA, Katz JS, Robinson JL. 2016. Delayed match-to-sample in working memory: A BrainMap meta-analysis. Biol Psychol. 120:10–20.

Edin F, Klingberg T, Johansson P, McNab F, Tegnér J, Compte A. 2009. Mechanism for top-down control of working memory capacity. Proc Natl Acad Sci U S A. 106:6802–6807.

Elton A, Gao W. 2014. Divergent task-dependent functional connectivity of executive control and salience networks. Cortex. 51:56–66.

Emch M, von Bastian CC, Koch K. 2019. Neural Correlates of Verbal Working Memory: An fMRI Meta-Analysis. Front Hum Neurosci. 13:180.

Emmert K, Kopel R, Sulzer J, Brühl AB, Berman BD, Linden DEJ, Horovitz SG, Breimhorst M, Caria A, Frank S, Johnston S, Long Z, Paret C, Robineau F, Veit R, Bartsch A, Beckmann CF, Van De Ville D, Haller S. 2016. Meta-analysis of real-time fMRI neurofeedback studies using individual participant data: How is brain regulation mediated? Neuroimage. 124:806–812.

Fishburn FA, Ludlum RS, Vaidya CJ, Medvedev AV. 2019. Temporal Derivative Distribution Repair (TDDR): A motion correction method for fNIRS. Neuroimage. 184:171–179.

Forbes NF, Carrick LA, McIntosh AM, Lawrie SM. 2009. Working memory in schizophrenia: a meta-analysis. Psychol Med. 39:889–905.

Friston KJ, Holmes AP, Worsley KJ, Poline JP, Frith CD, Frackowiak RS. 1994. Statistical parametric maps in functional imaging: a general linear approach. Human brain mapping. 2:189–210.

Garrison JR, Saviola F, Morgenroth E, Barker H, Lührs M, Simons JS, Fernyhough C, Allen P. 2021. Modulating medial prefrontal cortex activity using real-time fMRI neurofeedback: Effects on reality monitoring performance and associated functional connectivity. Neuroimage. 245:118640.

Goldway N, Jalon I, Keynan JN, Hellrung L, Horstmann A, Paret C, Hendler T. 2022. Feasibility and utility of amygdala neurofeedback. Neurosci Biobehav Rev. 138:104694.

Haller S, Kopel R, Jhooti P, Haas T, Scharnowski F, Lovblad KO, Scheffler K, Van De Ville D. 2013. Dynamic reconfiguration of human brain functional networks through neurofeedback. Neuroimage. 81:243–252.

Hawkinson JE, Ross AJ, Parthasarathy S, Scott DJ, Laramee EA, Posecion LJ, Rekshan WR, Sheau KE, Njaka ND, Bayley PJ, deCharms RC. 2012. Quantification of adverse events associated with functional MRI scanning and with real-time fMRI-based training. Int J Behav Med. 19:372–381.

Hernaus D, Casales Santa MM, Offermann JS, Van Amelsvoort T. 2017. Noradrenaline transporter blockade increases fronto-parietal functional connectivity relevant for working memory. Eur Neuropsychopharmacol. 27:399–410.

Hill AT, Fitzgerald PB, Hoy KE. 2016. Effects of Anodal Transcranial Direct Current Stimulation on Working Memory: A Systematic Review and Meta-Analysis of Findings From Healthy and Neuropsychiatric Populations. Brain Stimul. 9:197–208.

Horvath JC, Forte JD, Carter O. 2015. Quantitative Review Finds No Evidence of Cognitive Effects in Healthy Populations From Single-session Transcranial Direct Current Stimulation (tDCS). Brain Stimul. 8:535–550.

Hosseini SMH, Pritchard-Berman M, Sosa N, Ceja A, Kesler SR. 2016. Task-based neurofeedback training: A novel approach toward training executive functions. Neuroimage. 134:153–159.

Hou X, Xiao X, Gong Y, Li Z, Chen A, Zhu C. 2021. Functional Near-Infrared Spectroscopy Neurofeedback Enhances Human Spatial Memory. Front Hum Neurosci. 15:681193.

Hou X, Zhang Z, Zhao C, Duan L, Gong Y, Li Z, Zhu C. 2021. NIRS-KIT: a MATLAB toolbox for both restingstate and task fNIRS data analysis. Neurophotonics. 8:010802.

Huang J, Tan SP, Walsh SC, Spriggens LK, Neumann DL, Shum DH, Chan RC. 2014. Working memory dysfunctions predict social problem solving skills in schizophrenia. Psychiatry Res. 220:96–101.

Hudak J, Blume F, Dresler T, Haeussinger FB, Renner TJ, Fallgatter AJ, Gawrilow C, Ehlis AC. 2017. Near-Infrared Spectroscopy-Based Frontal Lobe Neurofeedback Integrated in Virtual Reality Modulates Brain and Behavior in Highly Impulsive Adults. Front Hum Neurosci. 11:425.

Hudak J, Rosenbaum D, Barth B, Fallgatter AJ, Ehlis AC. 2018. Functionally disconnected: A look at how study design influences neurofeedback data and mechanisms in attention-deficit/hyperactivity disorder. PLoS One. 13:e0200931.

Jahn H. 2013. Memory loss in Alzheimer’s disease. Dialogues Clin Neurosci. 15:445–454.

Jasper HH. 1958. Report of the committee on methods of clinical examination in electroencephalography. Electroencephalography and Clinical Neurophysiology. 10:370–375.

Jenkinson M, Bannister P, Brady M, Smith S. 2002. Improved optimization for the robust and accurate linear registration and motion correction of brain images. Neuroimage. 17:825–841.

Jolles DD, Grol MJ, Van Buchem MA, Rombouts SARB, Crone EA. 2010. Practice effects in the brain: Changes in cerebral activation after working memory practice depend on task demands. NeuroImage. 52:658–668.

Kaser M, Deakin JB, Michael A, Zapata C, Bansal R, Ryan D, Cormack F, Rowe JB, Sahakian BJ. 2017. Modafinil Improves Episodic Memory and Working Memory Cognition in Patients With Remitted Depression: A Double-Blind, Randomized, Placebo-Controlled Study. Biol Psychiatry Cogn Neurosci Neuroimaging. 2:115–122.

Kaufman AS. 1983. Test Review: Wechsler, D. Manual for the Wechsler Adult Intelligence Scale, Revised. New York: Psychological Corporation, 1981. Journal of Psychoeducational Assessment. 1:309–313.

Kimmig A-CS, Dresler T, Hudak J, Haeussinger FB, Wildgruber D, Fallgatter AJ, Ehlis A-C, Kreifelts B. 2019. Feasibility of NIRS-based neurofeedback training in social anxiety disorder: behavioral and neural correlates. Journal of Neural Transmission. 126:1175–1185.

Klingberg T. 2010. Training and plasticity of working memory. Trends Cogn Sci. 14:317–324.

Klugah-Brown B, Yu Y, Hu P, Agoalikum E, Liu C, Liu X, Yang X, Zeng Y, Zhou X, Yu X, Rypma B, Michael AM, Li X, Becker B, Biswal B. 2022. Effect of surgical mask on fMRI signals during task and rest. Communications Biology. 5:1004.

Kohl SH, Mehler DMA, Lührs M, Thibault RT, Konrad K, Sorger B. 2020. The Potential of Functional Near-Infrared Spectroscopy-Based Neurofeedback-A Systematic Review and Recommendations for Best Practice. Front Neurosci. 14:594.

Kohl SH, Veit R, Spetter MS, Günther A, Rina A, Lührs M, Birbaumer N, Preissl H, Hallschmid M. 2019. Real-time fMRI neurofeedback training to improve eating behavior by self-regulation of the dorsolateral prefrontal cortex: A randomized controlled trial in overweight and obese subjects. NeuroImage. 191:596–609.

Kumar S, Zomorrodi R, Ghazala Z, Goodman MS, Blumberger DM, Cheam A, Fischer C, Daskalakis ZJ, Mulsant BH, Pollock BG, Rajji TK. 2017. Extent of Dorsolateral Prefrontal Cortex Plasticity and Its Association With Working Memory in Patients With Alzheimer Disease. JAMA Psychiatry. 74:1266–1274.

Landau SM, Garavan H, Schumacher EH, D’Esposito M. 2007. Regional specificity and practice: dynamic changes in object and spatial working memory. Brain Res. 1180:78–89.

Landau SM, Schumacher EH, Garavan H, Druzgal TJ, D’Esposito M. 2004. A functional MRI study of the influence of practice on component processes of working memory. Neuroimage. 22:211–221.

Lee J, Park S. 2005. Working memory impairments in schizophrenia: a meta-analysis. J Abnorm Psychol. 114:599–611.

Li J, Yang X, Zhou F, Liu C, Wei Z, Xin F, Daumann B, Daumann J, Kendrick KM, Becker B. 2020. Modafinil enhances cognitive, but not emotional conflict processing via enhanced inferior frontal gyrus activation and its communication with the dorsomedial prefrontal cortex. Neuropsychopharmacology. 45:1026–1033.

Li K, Jiang Y, Gong Y, Zhao W, Zhao Z, Liu X, Kendrick KM, Zhu C, Becker B. 2019. Functional near-infrared spectroscopy-informed neurofeedback: regional-specific modulation of lateral orbitofrontal activation and cognitive flexibility. Neurophotonics. 6:025011.

Linden DE. 2007. The working memory networks of the human brain. Neuroscientist. 13:257–267.

Liu N, Cui X, Bryant DM, Glover GH, Reiss AL. 2015. Inferring deep-brain activity from cortical activity using functional near-infrared spectroscopy. Biomed Opt Express. 6:1074–1089.

Mancuso LE, Ilieva IP, Hamilton RH, Farah MJ. 2016. Does Transcranial Direct Current Stimulation Improve Healthy Working Memory?: A Meta-analytic Review. J Cogn Neurosci. 28:1063–1089.

Marx AM, Ehlis AC, Furdea A, Holtmann M, Banaschewski T, Brandeis D, Rothenberger A, Gevensleben H, Freitag CM, Fuchsenberger Y, Fallgatter AJ, Strehl U. 2014. Near-infrared spectroscopy (NIRS) neurofeedback as a treatment for children with attention deficit hyperactivity disorder (ADHD)-a pilot study. Front Hum Neurosci. 8:1038.

McKendrick R, Ayaz H, Olmstead R, Parasuraman R. 2014. Enhancing dual-task performance with verbal and spatial working memory training: Continuous monitoring of cerebral hemodynamics with NIRS. NeuroImage. 85:1014–1026.

McLaren DG, Ries ML, Xu G, Johnson SC. 2012. A generalized form of context-dependent psychophysiological interactions (gPPI): A comparison to standard approaches. NeuroImage. 61:1277–1286.

Molnar-Szakacs I, Uddin LQ. 2022. Anterior insula as a gatekeeper of executive control. Neurosci Biobehav Rev. 139:104736.

Nikolin S, Tan YY, Schwaab A, Moffa A, Loo CK, Martin D. 2021. An investigation of working memory deficits in depression using the n-back task: A systematic review and meta-analysis. Journal of Affective Disorders. 284:1–8.

Nouchi R, Nouchi H, Dinet J, Kawashima R. 2021. Cognitive Training with Neurofeedback Using NIRS Improved Cognitive Functions in Young Adults: Evidence from a Randomized Controlled Trial. Brain Sci. 12.

Oda Y, Sato T, Nambu I, Wada Y. 2018. Real-Time Reduction of Task-Related Scalp-Hemodynamics Artifact in Functional Near-Infrared Spectroscopy with Sliding-Window Analysis. Applied Sciences. 8:149.

Olesen PJ, Westerberg H, Klingberg T. 2004. Increased prefrontal and parietal activity after training of working memory. Nat Neurosci. 7:75–79.

Pamplona GSP, Heldner J, Langner R, Koush Y, Michels L, Ionta S, Scharnowski F, Salmon CEG. 2020. Networkbased fMRI-neurofeedback training of sustained attention. Neuroimage. 221:117194.

Pinti P, Tachtsidis I, Hamilton A, Hirsch J, Aichelburg C, Gilbert S, Burgess PW. 2020. The present and future use of functional near-infrared spectroscopy (fNIRS) for cognitive neuroscience. Ann N Y Acad Sci. 1464:5–29.

Ramos AA, Machado L. 2021. A Comprehensive Meta-analysis on Short-term and Working Memory Dysfunction in Parkinson’s Disease. Neuropsychol Rev. 31:288–311.

Rance M, Walsh C, Sukhodolsky DG, Pittman B, Qiu M, Kichuk SA, Wasylink S, Koller WN, Bloch M, Gruner P, Scheinost D, Pittenger C, Hampson M. 2018. Time course of clinical change following neurofeedback. Neuroimage. 181:807–813.

Rasetti R, Mattay VS, Stankevich B, Skjei K, Blasi G, Sambataro F, Arrillaga-Romany IC, Goldberg TE, Callicott JH, Apud JA, Weinberger DR. 2010. Modulatory effects of modafinil on neural circuits regulating emotion and cognition. Neuropsychopharmacology. 35:2101–2109.

Salmi J, Nyberg L, Laine M. 2018. Working memory training mostly engages general-purpose large-scale networks for learning. Neuroscience & Biobehavioral Reviews. 93:108–122.

Sayala S, Sala JB, Courtney SM. 2005. Increased Neural Efficiency with Repeated Performance of a Working Memory Task is Information-type Dependent. Cerebral Cortex. 16:609–617.

Scharnowski F, Weiskopf N. 2015. Cognitive enhancement through real-time fMRI neurofeedback. Current Opinion in Behavioral Sciences. 4:122–127.

Schommartz I, Dix A, Passow S, Li SC. 2020. Functional Effects of Bilateral Dorsolateral Prefrontal Cortex Modulation During Sequential Decision-Making: A Functional Near-Infrared Spectroscopy Study With Offline Transcranial Direct Current Stimulation. Front Hum Neurosci. 14:605190.

Shen J, Zhang G, Yao L, Zhao X. 2015. Real-time fMRI training-induced changes in regional connectivity mediating verbal working memory behavioral performance. Neuroscience. 289:144–152.

Sherwood MS, Kane JH, Weisend MP, Parker JG. 2016. Enhanced control of dorsolateral prefrontal cortex neurophysiology with real-time functional magnetic resonance imaging (rt-fMRI) neurofeedback training and working memory practice. Neuroimage. 124:214–223.

Sitaram R, Ros T, Stoeckel L, Haller S, Scharnowski F, Lewis-Peacock J, Weiskopf N, Blefari ML, Rana M, Oblak E, Birbaumer N, Sulzer J. 2017. Closed-loop brain training: the science of neurofeedback. Nat Rev Neurosci. 18:86–100.

Sligte IG, Wokke ME, Tesselaar JP, Steven Scholte H, Lamme VAF. 2011. Magnetic stimulation of the dorsolateral prefrontal cortex dissociates fragile visual short-term memory from visual working memory. Neuropsychologia. 49:1578–1588.

Smith J, Browning M, Conen S, Smallman R, Buchbjerg J, Larsen KG, Olsen CK, Christensen SR, Dawson GR, Deakin JF, Hawkins P, Morris R, Goodwin G, Harmer CJ. 2018. Vortioxetine reduces BOLD signal during performance of the N-back working memory task: a randomised neuroimaging trial in remitted depressed patients and healthy controls. Mol Psychiatry. 23:1127–1133.

Soekadar SR, Kohl SH, Mihara M, von Lühmann A. 2021. Optical brain imaging and its application to neurofeedback. Neuroimage Clin. 30:102577.

Spielberger CD. 1970. Manual for the state-trait anxietry, inventory. Consulting Psychologist.

Sun X, Yue S, Duan M, Yao D, Luo C. 2023. Psychosocial intervention for schizophrenia. Brain-Apparatus Communication: A Journal of Bacomics. 1–11.

Tang Y, Chen Z, Jiang Y, Zhu C, Chen A. 2021. From reversal to normal: Robust improvement in conflict adaptation through real-time functional near infrared spectroscopy-based neurofeedback training. Neuropsychologia. 157:107866.

Teixeira-Santos AC, Moreira CS, Magalhães R, Magalhães C, Pereira DR, Leite J, Carvalho S, Sampaio A. 2019. Reviewing working memory training gains in healthy older adults: A meta-analytic review of transfer for cognitive outcomes. Neurosci Biobehav Rev. 103:163–177.

Thibault RT, MacPherson A, Lifshitz M, Roth RR, Raz A. 2018. Neurofeedback with fMRI: A critical systematic review. Neuroimage. 172:786–807.

Wager TD, Smith EE. 2003. Neuroimaging studies of working memory: a meta-analysis. Cogn Affect Behav Neurosci. 3:255–274.

Wang H, He W, Wu J, Zhang J, Jin Z, Li L. 2019. A coordinate-based meta-analysis of the n-back working memory paradigm using activation likelihood estimation. Brain Cogn. 132:1–12.

Wang X-L, Du M-Y, Chen T-L, Chen Z-Q, Huang X-Q, Luo Y, Zhao Y-J, Kumar P, Gong Q-Y. 2015. Neural correlates during working memory processing in major depressive disorder. Progress in Neuro-Psychopharmacology and Biological Psychiatry. 56:101–108.

Watanabe T, Sasaki Y, Shibata K, Kawato M. 2017. Advances in fMRI Real-Time Neurofeedback. Trends in Cognitive Sciences. 21:997–1010.

Watson D, Clark LA, Tellegen A. 1988. Development and validation of brief measures of positive and negative affect: the PANAS scales. J Pers Soc Psychol. 54:1063–1070.

Weiss F, Zhang J, Aslan A, Kirsch P, Gerchen MF. 2022. Feasibility of training the dorsolateral prefrontal-striatal network by real-time fMRI neurofeedback. Scientific Reports. 12:1669.

Westerberg H, Klingberg T. 2007. Changes in cortical activity after training of working memory—a single-subject analysis. Physiology & Behavior. 92:186–192.

Wischnewski M, Mantell KE, Opitz A. 2021. Identifying regions in prefrontal cortex related to working memory improvement: A novel meta-analytic method using electric field modeling. Neurosci Biobehav Rev. 130:147–161.

Wu W-J, Cui L-B, Cai M, Peng Z-W, Zhang W-C, Lv S, Xu J-Y, Hu Y, Li G, von Deneen KM, Zhu C-Z, Wang H-N, Zhang Y. 2022. A parallel-group study of near-infrared spectroscopy-neurofeedback in children with attention deficit hyperactivity disorder. Psychiatry Research. 309:114364.

Xu X, Xin F, Liu C, Chen Y, Yao S, Zhou X, Zhou F, Huang Y, Dai J, Wang J, Zou Z, Kendrick KM, Zhou B, Becker B. 2022. Disorder-and cognitive demand-specific neurofunctional alterations during social emotional working memory in generalized anxiety disorder and major depressive disorder. J Affect Disord. 308:98–105.

Yamada T, Umeyama S, Matsuda K. 2012. Separation of fNIRS signals into functional and systemic components based on differences in hemodynamic modalities. PLoS One. 7:e50271.

Yamamoto T, Kato T. 2002. Paradoxical correlation between signal in functional magnetic resonance imaging and deoxygenated haemoglobin content in capillaries: a new theoretical explanation. Phys Med Biol. 47:1121–1141.

Yao S, Becker B, Geng Y, Zhao Z, Xu X, Zhao W, Ren P, Kendrick KM. 2016. Voluntary control of anterior insula and its functional connections is feedback-independent and increases pain empathy. Neuroimage. 130:230–240.

Yu L, Long Q, Tang Y, Yin S, Chen Z, Zhu C, Chen A. 2021. Improving Emotion Regulation Through Real-Time Neurofeedback Training on the Right Dorsolateral Prefrontal Cortex: Evidence From Behavioral and Brain Network Analyses. Front Hum Neurosci. 15:620342.

Zhang G, Yao L, Shen J, Yang Y, Zhao X. 2015. Reorganization of functional brain networks mediates the improvement of cognitive performance following real-time neurofeedback training of working memory. Hum Brain Mapp. 36:1705–1715.

Zhang G, Yao L, Zhang H, Long Z, Zhao X. 2013. Improved working memory performance through self-regulation of dorsal lateral prefrontal cortex activation using real-time fMRI. PLoS One. 8:e73735.

Zhang Q, Zhang G, Yao L, Zhao X. 2015. Impact of real-time fMRI working memory feedback training on the interactions between three core brain networks. Front Behav Neurosci. 9:244.

Zhao W, Becker B, Yao S, Ma X, Kou J, Kendrick KM. 2019. Oxytocin Enhancement of the Placebo Effect May Be a Novel Therapy for Working Memory Impairments. Psychother Psychosom. 88:125–126.

Zhao Z, Yao S, Li K, Sindermann C, Zhou F, Zhao W, Li J, Lührs M, Goebel R, Kendrick KM, Becker B. 2019. Real-Time Functional Connectivity-Informed Neurofeedback of Amygdala-Frontal Pathways Reduces Anxiety. Psychother Psychosom. 88:5–15.

Zweerings J, Sarkheil P, Keller M, Dyck M, Klasen M, Becker B, Gaebler AJ, Ibrahim CN, Turetsky BI, Zvyagintsev M, Flatten G, Mathiak K. 2020. Rt-fMRI neurofeedback-guided cognitive reappraisal training modulates amygdala responsivity in posttraumatic stress disorder. Neuroimage Clin. 28:102483.

