## Supplemental Material for "A brief real-time fNIRS-informed neurofeedback training of the prefrontal cortex changes brain activity and connectivity during subsequent working memory challenge"

### Supplementary Materials

Xi Yang et al.,

Contact

Benjamin Becker,; Yihan Jiang,

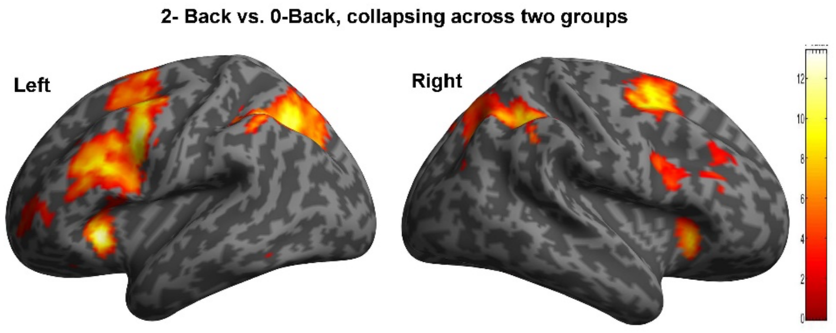


Supplementary Figure 1: Brain Activations Related to WM load. The figure shows regions of significant activation for the 2-back vs 0-back contrast ($P_{FWE\text{-}corrected} < 0.05$).

| Region/cluster | Peak MNI coordinates | | | Cluster size in voxels (k) | Peak T score | $P_{FWE\text{-}corrected}$ < 0.05 |
|---|---|---|---|---|---|---|
| | X | Y | Z | | | |
| | 2-back > 0-back contrast (k>100) | | | | | |
| Medial Frontal Gyrus | -8 | 12 | 50 | 2741 | 13.94 | <0.001 |
| Left Inferior Frontal Gyrus/insula | -28 | 24 | -2 | 395 | 13.32 | <0.001 |
| Left Inferior Frontal Gyrus/Precentral | -44 | 4 | 36 | 1224 | 12.70 | <0.001 |
| Left Parietal lobe | -32 | -54 | 38 | 1038 | 13.27 | <0.001 |
| Left Precuneus | -12 | -68 | 46 | 376 | 8.96 | <0.001 |
| Right Inferior Parietal Lobule | 36 | -46 | 40 | 640 | 10.48 | <0.001 |
| Left Precuneus | 12 | -68 | 44 | 276 | 8.27 | <0.001 |
| Right Insula | 28 | 26 | 0 | 142 | 9.73 | <0.001 |
| Right Cerebellum | 26 | -62 | -38 | 313 | 8.15 | <0.001 |
| Right Inferior Frontal Gyrus | 46 | 6 | 24 | 178 | 7.55 | <0.001 |
| Right Middle Frontal Gyrus | 38 | 32 | 26 | 260 | 6.48 | <0.001 |

Supplementary Table 1: Brain activations related to WM load
